# Controlled Self-Assembly of λ-DNA Networks with the Synergistic Effect of DC Electric Field

**DOI:** 10.1101/774901

**Authors:** M. Gao, J. Hu, Y. Wang, M. Liu, J. Wang, Z. Song, H. Xu, C. Hu, Z. Wang

**Affiliations:** International Research Centre for Nano Handling and Manufacturing of China, Changchun University of Science and Technology; Changchun University of Science and Technology

**Keywords:** Atomic force microscopy, self-assembly, DNA network, DC electric field

## Abstract

Large-scale and morphologically controlled self-assembled λ-DNA networks were successfully constructed by the synergistic effect of DC electric field. The effect of DNA concentration, direction and intensity of the electric field, even the modification of the mica surface using Mg^2+^ on the characteristics of the as-prepared DNA networks were investigated in detail by atomic force microscopy (AFM). It was found that the horizontal electric field was more advantageous to the formation of DNA networks with more regular structures. At the same concentration, the height of DNA network was not affected significantly by the intensity change of the horizontal electric field. The modification of Mg^2+^ on mica surface increased the aggregation of DNA molecules, which contributed to the morphological change of the DNA networks. Furthermore, DNA molecules were obviously stretched in both horizontal and vertical electric fields at low DNA concentrations.

**Statement of significance:** Through the synergistic effect of DC electric field, a series of large-scale and morphologically controlled self-assembled λ-DNA networks were successfully fabricated. We found that the horizontal electric field was more advantageous to the formation of DNA networks with more regular structures. At the same concentration of DNA solution, the height of DNA network was not affected significantly by the intensity change of the horizontal electric field. The modification of Mg^2+^ on mica surface increased the aggregation of DNA molecules, which contributed to the morphological change of the DNA networks. We suggest this study will promote the understanding on the preparation of controllable self-assembled λ-DNA networks and the application of DNA networks.

## Introduction

As a promising biomolecule, DNA not only plays an indisputable role in life science and bioengineering, but also attracts extensive attention in the field of nanotechnology (1). The intrinsic ability of DNA to arrange itself has promoted the studies of surface assembly properties of DNA molecules (2). Self-assembly enables DNA molecules to be combined into complex structures in a relatively simple step, thus providing remarkably flexible way in the design of various molecular patterns (3–9), which have potential applications in medicine and engineering as molecular functional devices (10–14), DNA-based electrochemical biosensors, or act as a scaffold to construct nanostructures (15), such as nanoparticles (16–19), nanorods (20–22), nanowires (23–25) and two-dimensional nanoparticle networks (26). However, it is still a challenge to realize the controllable assembly of DNA molecules into highly oriented interlaced patterns on a large scale, thus makes it an obstacle for their future application in nanotechnology.

Over the past decades, research into exploring simple and effective methods for controllable DNA self-assembly has never ceased. Among them, the preparation and formation mechanism of DNA self-assembled networks with unique two-dimensional and even three-dimensional structures on a solid surface have attracted wide attention. DNA networks formed by self-assembly on solid surfaces were generally prepared by facial deposition method using DNA solution with relatively high concentration. The density, height and the coverage of DNA networks were usually regulated as reported by adjusting the concentration of DNA solution, deposition time and the type of substrates including mica (27–28), silicon (29), gold (30) or other materials (31–32). In addition, whether the surface was modified or not by salt cations was also a neglectable factor for the fabrication of regular networks (33). Nevertheless, although large-scale two-dimensional DNA networks could be obtained through the methods above, there are still restrictions in the structural diversity of the prepared DNA networks. Furthermore, it has been proved that DNA molecules exhibit a tendency of movement and stretching (34) under electric fields, however, the self-assembly of DNA molecules into specific patterns under an electric field has rarely been reported.

In this work, a series of large-scale and morphologically controlled self-assembled λ-DNA networks were successfully fabricated by the synergistic effect of DC electric field. The direction of the DC electric field was divided into horizontal electric field (HEF) and vertical electric field (VEF), and the intensity of the electric field was simply adjusted by changing the voltage. Different influencing factors including the DNA concentration, direction and intensity of the electric field, and even the modification using Mg^2+^ for the mica surface were investigated by atomic force microscopy (AFM) in detail on the morphology of the as-prepared DNA networks. The obtained information will promote the understanding on the preparation of controllable self-assembled λ-DNA networks and the application of DNA networks.

## Experimental sections

### Materials

All chemicals were used as received without purification throughout the study. λ-DNA (48502 bp) was purchased from Thermo Fisher Scientific Company (China). MgCl_2_·6H_2_O was obtained from Beijing Chemical Works (China). Ultrapure water used during the experiments (≥18.2 MΩ cm) was purified with a Milli-Q water purification system. The 1.0×1.0 cm^2^ fluorphlogopite (KMg_3_(AlSi_3_O10)F_2_) mica square pieces were purchased from Taiyuan Fluorphlogopite Mica Company (China). The mica used before the subsequent experiments was freshly cleaved.

### DNA solution preparation

The 500 ng/μL stock solution of λ-DNA was diluted to different concentrations with ultrapure water to the concentrations of 1 ng/μL, 5 ng/μL and 10 ng/μL for sample preparations.

### DNA sample preparation under DC electric fields

In order to investigate the effect of electric field direction on the formation of DNA samples in detail, two forms of electric fields including the horizontal electric field (HEF) and the vertical electric field (VEF) were prepared respectively as shown in Fig. 1. Herein, the E363A instrument from the Keysight Technologies Company (USA) was used as the DC electric source. A pair of indium-tin oxide (ITO) glasses (2.0×2.0 cm^2^, 100 Ω/cm^2^) with the ITO layer thickness of 230Å±50Å were used as the electrodes. The examined distance between the two electrodes of the horizontal and vertical electric fields was 10 mm and 2 mm, respectively, and the mica was placed between the electrodes. After applying the DC voltage for a few seconds, 3 μL of DNA solution was dropped onto the center of mica surface and the DC voltage was then remained for 7 min. Ultimately, the sample was dried in air and prepared for AFM imaging.

**Fig. 1.**
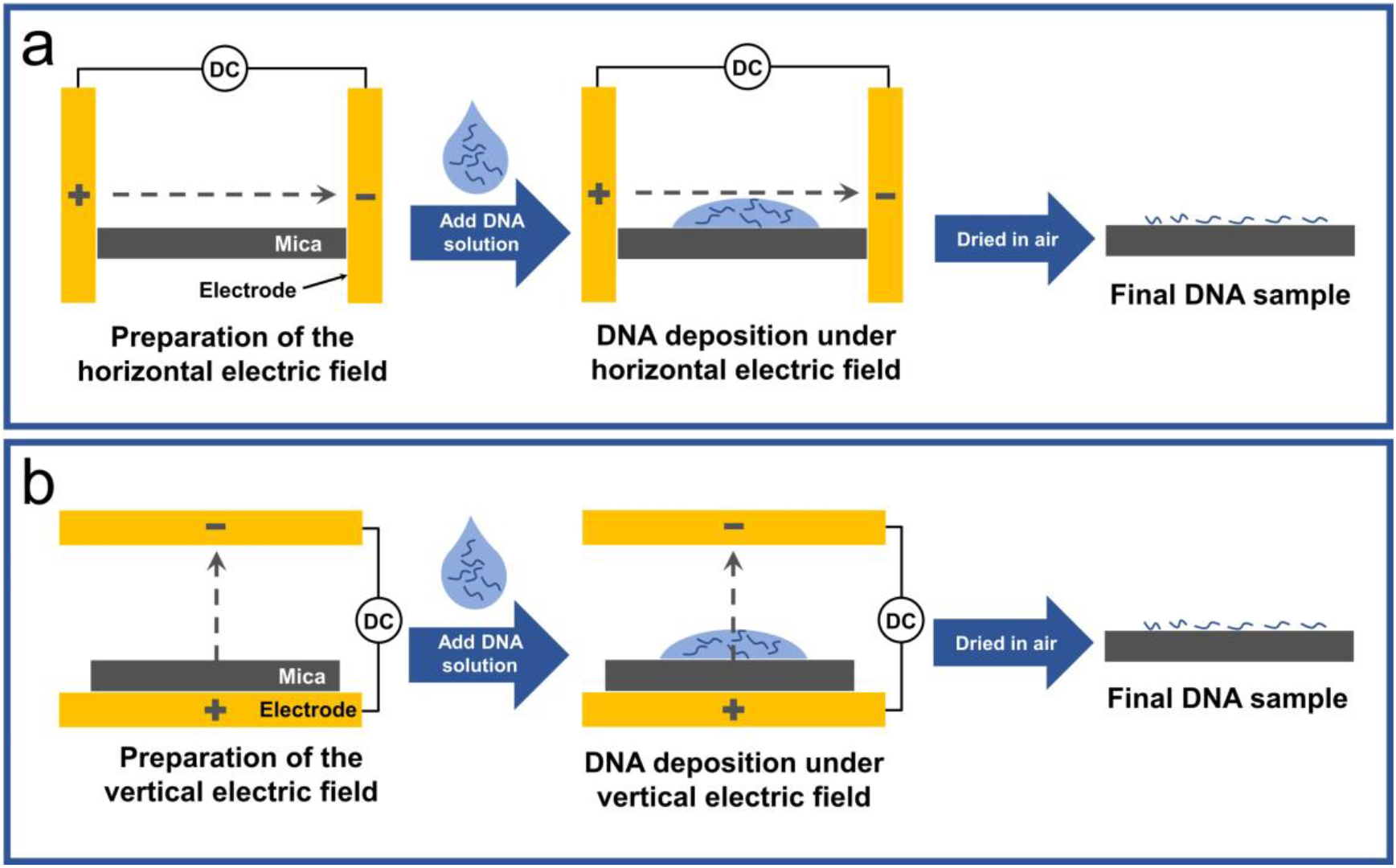
Schematic drawing for the preparation of DNA samples under horizontal (a) and vertical (b) electric fields.

To figure out the effect of Mg^2+^ on the deposition of DNA under the DC electric field, a Mg^2+^ modified mica surface was prepared as follows. 10 mM MgCl_2_ aqueous solution was prepared by dissolving 0.2033 g MgCl_2_·6H_2_O into 100 mL ultrapure water. Then a droplet of the MgCl_2_ aqueous solution (2.5 μL) was dropped onto a freshly cleaved mica surface and being spread out. After remained for 3 min, the mica was rinsed with ultrapure water and blown dry with compressed air before dropping the DNA solution. The Drop Shape Analyzer-DSA100 (Germany) was utilized to analyze the contact angle of the mica before and after Mg^2+^ modifications.

In a control experiment, the DNA samples adsorbed on bare and Mg^2+^ modified mica surfaces without DC electric fields were also prepared according to the same procedures respectively as mentioned above.

It should be noted that for the vertical electric field, although not shown in this figure, the downward direction of the electric field was also examined in this experiment.

### Atomic force microscopy measurements

The AFM experiments were carried out using the Agilent 5500 scanning probe microscope (SPM) from Agilent Technologies Company (USA). The AFM images were acquired in tapping mode using Tap300Al-G AFM probes (spring constant, 40 N/m) below the resonant frequency (typically, 200 ~ 400 KHz) under ambient conditions at room temperature.

## Results and discussion

As a control experiment in subsequent studies of DNA deposition under electric fields, various concentrations of DNA deposited on bare mica without any electric field were prepared. It should be noted that all the AFM images used in this paper were typical of those obtained from at least five separated samples. It was clear to see the changes of DNA molecular assembly due to the increase of concentration as shown in Fig. 2. In the low DNA concentration of 1 ng/μL (Fig. 2a), the DNA molecules dispersed irregularly with slight intermolecular entanglement. When the DNA concentration increased to 5 ng/μL (Fig. 2b), the interconnection degree between DNA molecules increased, which led to the tendency of forming a DNA molecular network. Nevertheless, at this moment, the size of meshes was still large and nonuniform with prominent knots, and the end of the free DNA chain could be seen at the edge of the mesh. When the concentration of DNA reached 10 ng/μL as shown in Fig. 2c, an almost complete DNA network with approximate polygonal meshes appeared in the whole field of vision. The DNA network meshes became more uniform and smaller in size, clearer in edge without redundant bifurcation. Since the single chain height of double stranded DNA (dsDNA) measured by AFM in air was reported to be about 0.5 nm (35), the fibers of the DNA network were measured to be predominantly 8~16 dsDNA high according to the height profile corresponded with the AFM image, which was higher than the DNA fibers in Fig. 2a and Fig. 2b. Therefore, the fibers in a DNA network might be DNA bundles formed by overlapping or interlacing multiple dsDNA chains (36).

**Fig. 2.**
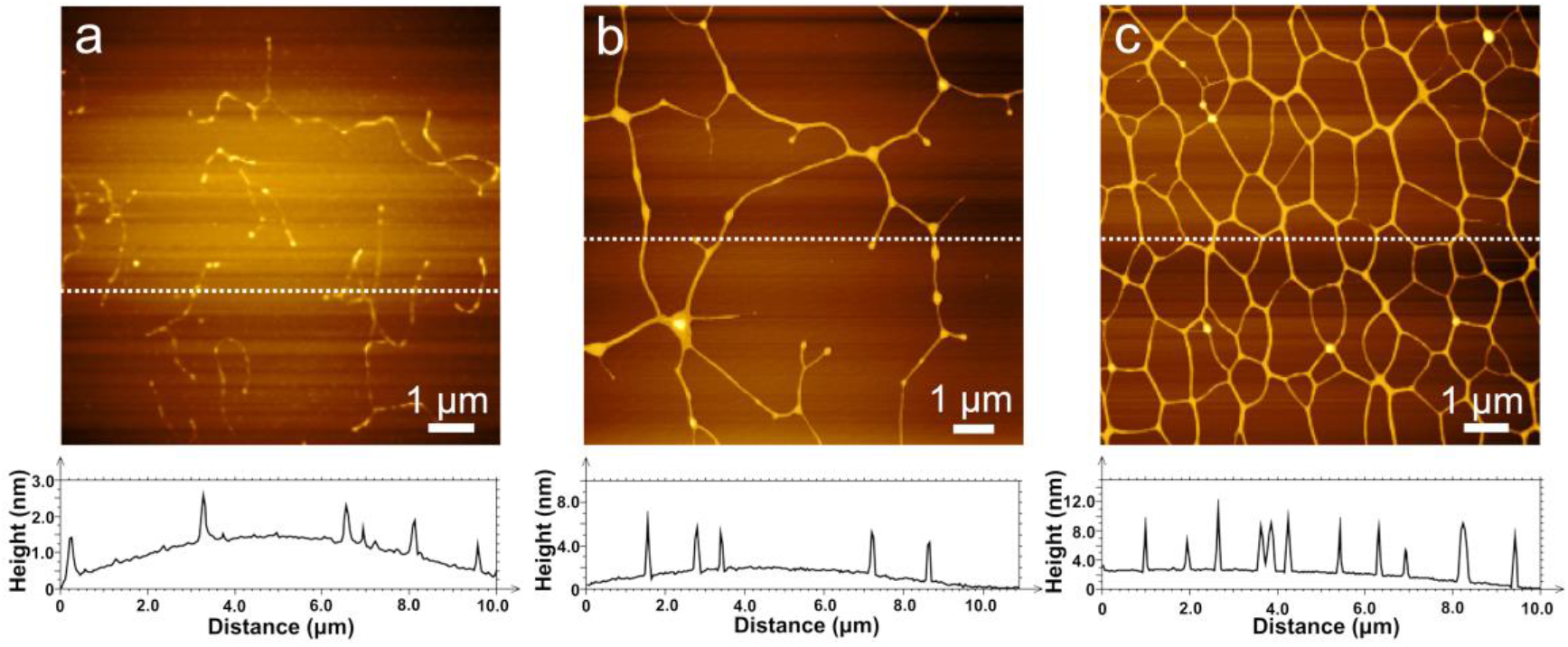
Typical AFM images (upper part) and height profiles (lower part) of DNA on the bare mica prepared with the different concentrations of DNA solution: 1 ng/μL (a), 5 ng/μL(b) and 10 ng/μL (c).

Fig. 3 shows the deposition of 1 ng/μL DNA on the bare mica under horizontal electric fields which are parallel to the mica surface at different voltages. Herein, the positive and negative electrodes are noted as + and −, respectively. The middle part between the positive and negative electrodes was selected to obtain the AFM images. Compared with the case without applying the electric field (Fig. 2a), at the same concentration of DNA solution, the number of DNA fibers in the field of vision increased significantly even at a low voltage of 0.5 V as shown in Fig. 3a. In addition, the strands of DNA molecules partially contacted and twined around each other to form the network, which was relatively relaxed and roughly extended along the direction of the horizontal electric field. This might be attributed to the fact that the electric field force increased the contact and crossover opportunities of the initially free-distributed DNA molecules in the solution and extended the network in a certain direction. At the voltage of 1.0 V (Fig. 3b), the density of DNA fibers in the field of vision decreased and the extension degree of the DNA molecules increased. Moreover, the extension direction of the whole fibers was deflected for a certain angle in comparison with that of Fig. 3a. When the voltage increased to a higher value of 2.0 V (Fig. 3c), the DNA molecules were further stretched, where the arranged direction of DNA fibers was identical and regularly deviated for a larger angle. Moreover, it was interesting that the increase of electric field intensity did not cause significant changes in the height of DNA fibers.

**Fig. 3.**
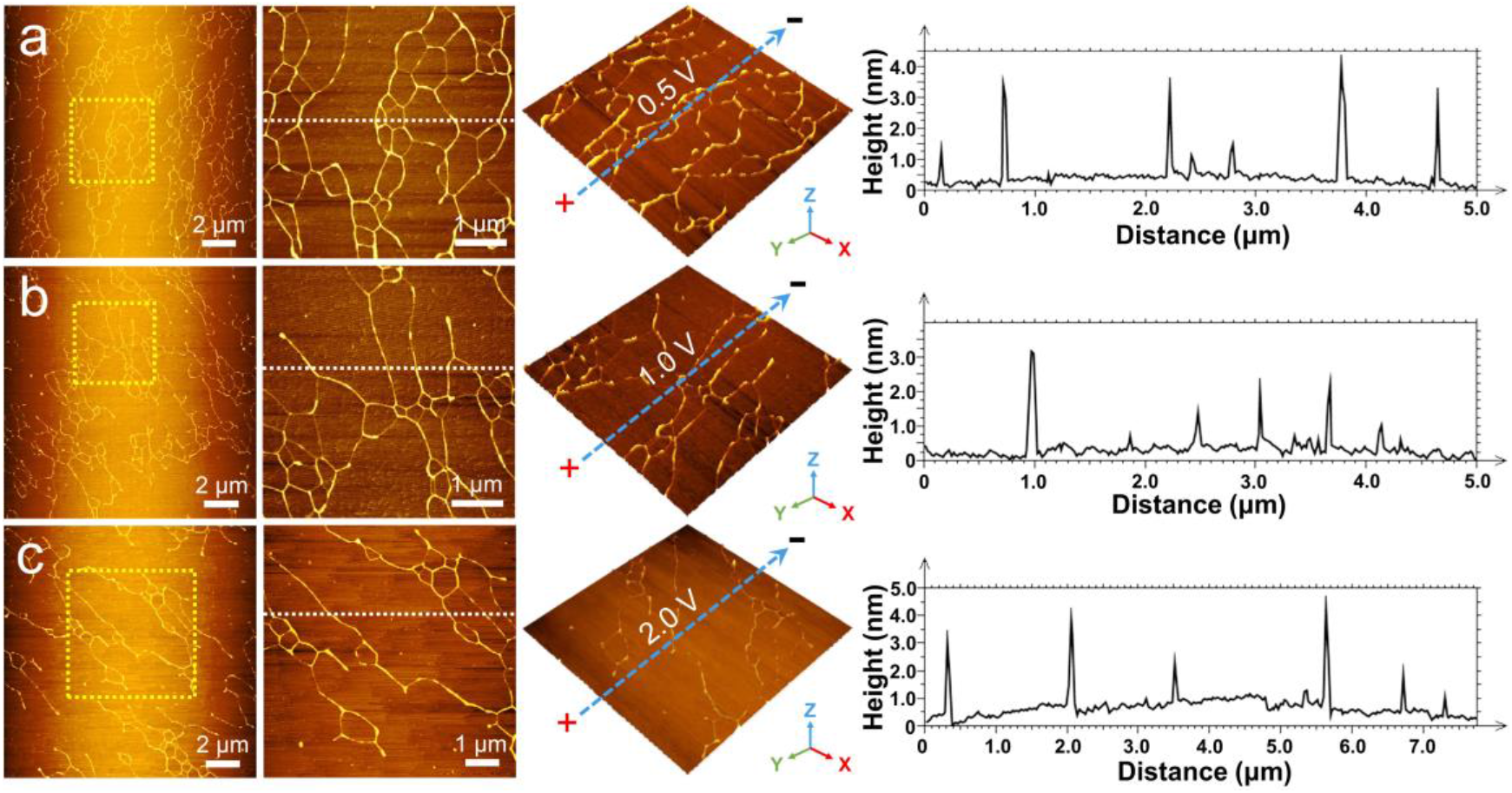
Typical AFM images (the first column), local amplification images (the second column) and the corresponding 3D topographic maps (the third column) of 1 ng/μL DNA on the bare mica produced under the different HEF DC voltages: 0.5 V (a), 1.0 V (b) and 2.0 V (c). (Note: The blue dotted arrow on the 3D topographic maps represents the direction of the horizontal electric field.) The fourth column is the height profiles of the transversal section marked with the white dashed line in the local amplification images.

Since the concentration of DNA solution was confirmed to be a crucial factor of forming the DNA network (37) and the horizontal electric field force did affect the deposition of DNA molecules from the above results, the self-assembly rule of DNA with higher concentration (10 ng/μL) under the horizontal electric field at different voltages was further investigated as shown in Fig. 4. The selection of the AFM imaging region was the same as that of Fig. 3. When the concentration of DNA increased, the DNA molecules were assembled into arrays completely different from those in Fig. 3a under the same voltage of electric field as shown in Fig. 4a. The arrays assembled was herringbone-like with forked tails and oriented toward the same direction, which was almost perpendicular to the direction of electric field. The periodicity of the arrangement was obviously reflected by the bright lines passing through the central point in the Fourier transform image. In addition to the alteration of arrangement, another obvious phenomenon was that the width of the herringbone fibers assembled by DNA molecules increased without significant change of the height. This might be due to the DNA bundles overlapped or intertwined with multiple dsDNA chains horizontally accumulated into wider rather than thicker fibers under the horizontal electric field, which could also be observed from the local amplification image and the height profiles.

**Fig. 4.**
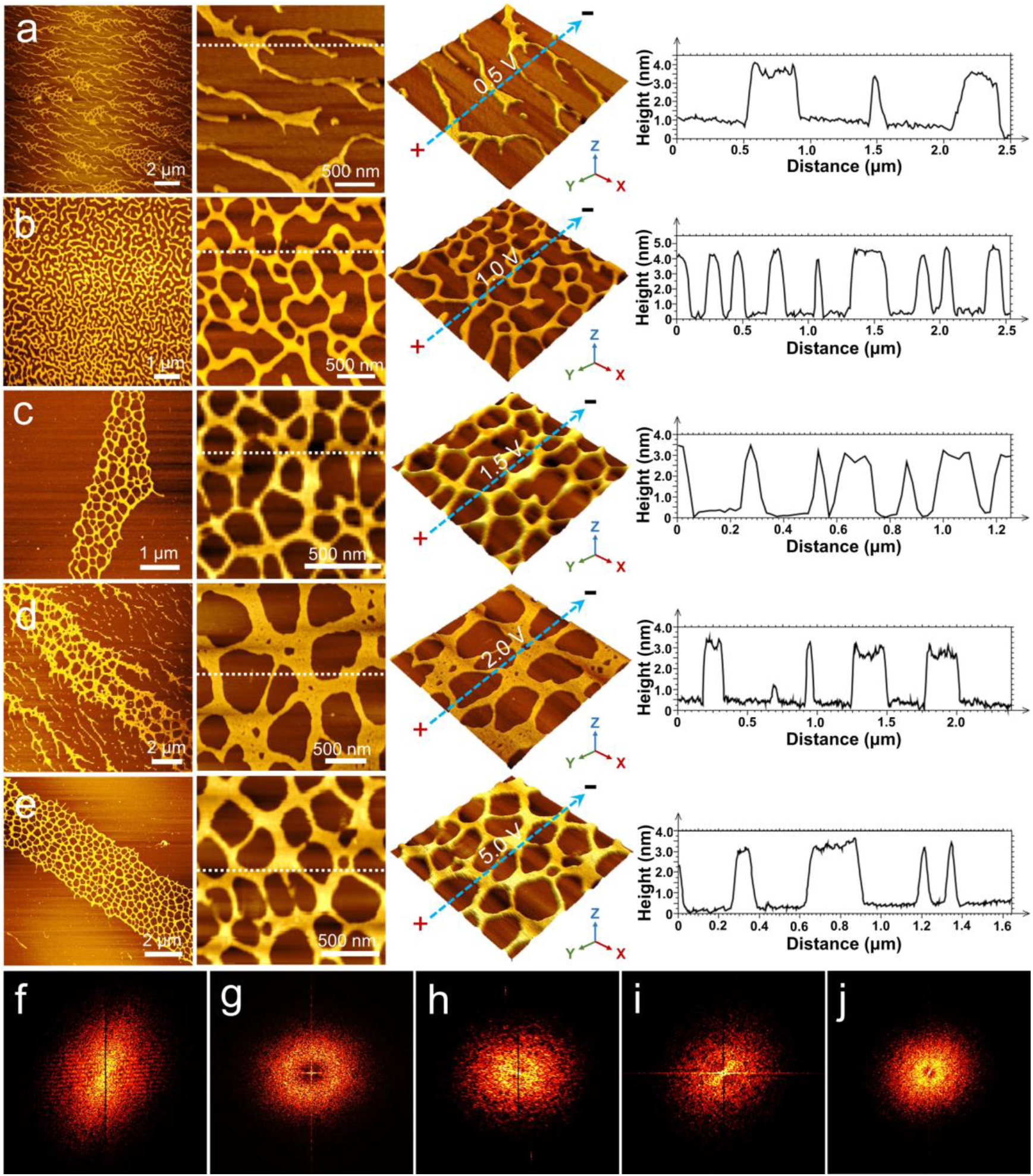
Typical AFM images (the first column), local amplification images (the second column) and the corresponding 3D topographic maps (the third column) of 10 ng/μL DNA on the bare mica produced under the different HEF DC voltages: 0.5 V (a), 1.0 V (b), 1.5 V (c), 2.0 (d) and 5.0 V (e). (Note: The blue dotted arrow on the 3D topographic maps represents the direction of the horizontal electric field.) The fourth column is the height profiles of the transversal section marked with the white dashed line in the local amplification images. (f)~(j) are corresponding Fourier transform images of (a)~(e) the first column typical AFM images.

Different from the whole stretching of DNA network at a low concentration after the electric field enhancement, the increase of the voltage from 0.5 V to 1.0 V accompanied with a high DNA concentration led to the formation of a large-scale network with unique maze-like structures, uniformly distributed on the surface of mica as shown in Fig. 4b. The uniform distribution of the network can be further confirmed by the nearly circular diffraction ring in the Fourier transform image (Fig. 4g). The meshes in this network were different in size without the evident directionality, the edges were smooth without obvious corners. In addition, although the coverage and density of the DNA network, even the width of DNA fibers formed in this way were much higher than those of the network shown in Fig. 2c, the height of the fibers did not have significant change.

When the voltage increased to 1.5 V, the DNA network arrays with the parallel arrangement appeared on the surface of the bare mica, which were extended along the direction of the electric field as shown in Fig. 4c. Besides, the direction of periodic arrangement can also be reflected from the direction of bright lines passing through the central point in Fig. 4h. The meshes were uniform in size with the polygonal shape and closely connected with each other. The width and height of the fibers were similar to those in Fig. 4b. The horizontal accumulation of the DNA bundles increased to a higher level as the voltage increased to 2.0 V, where the thin layer with obvious micropores formed by the intermolecular accumulation were clearly visible as shown in the local amplification image of Fig. 4d. The thin layers were connected and assembled into a DNA network with seemingly lacerated boundaries, and the herringbone fibers were parallelly distributed to the side of the network possibly formed due to the transverse pulling effect of electric field force. It was worth noting that the height of the DNA network formed at this voltage still had no significant change with the increase of the voltage.

While as the voltage rose to 5 V, the DNA network arrays in the parallel arrangement with clear and regular boundaries appeared again on the surface of the bare mica, which were displayed distinctly in Fig. 4e. Compared with the DNA network arrays shown in Fig. 4d, the shape of the arrays and the distribution of the meshes were more regular, which could be seen from the highly bright near-circular diffraction ring in Fig. 4j. The arrangement periodicity of the arrays was much clearer since there was a narrower bright line passing through the central point than that in Fig. 4i. Furthermore, similar to Fig. 4d, the network was fabricated by the connected and assembled thin layers and there was no significant change in the height of the network. Therefore, by adjusting the intensity of electric field and the concentration of DNA solution, it is possible to fabricate network arrays with uniform heights and different morphologies.

In order to figure out the influence of surface properties on DNA self-assembly under the DC electric field, the mica modified by Mg^2+^ was prepared for the further examination. Fig. 5a is the AFM result of a control experiment where the sample was not treated by the electric field. The DNA chains were loosely and disorderly distributed on the surface of mica without obvious intermolecular contact, crossover and overlap for the formation of regular networks. Fig. 5b displays the sample treated by the HEF DC voltage of 0.5 V on Mg^2+^ modified mica. The modification of Mg^2+^ certainly affected the DNA aggregation behavior that a network with large meshes appeared in the image, which was quite different with the network shown in Fig. 4b at the same voltage. In addition to morphological changes, the height of DNA networks has also increased to a certain degree. As the voltage increased to 1.0 V, the DNA molecules accumulated with each other severely to form parallel long arrays, in which the height of the arrays even reached nearly 100 nm. Therefore, the surface modification of mica should be an important factor for the influence of DNA self-assembly in the electric field.

**Fig. 5.**
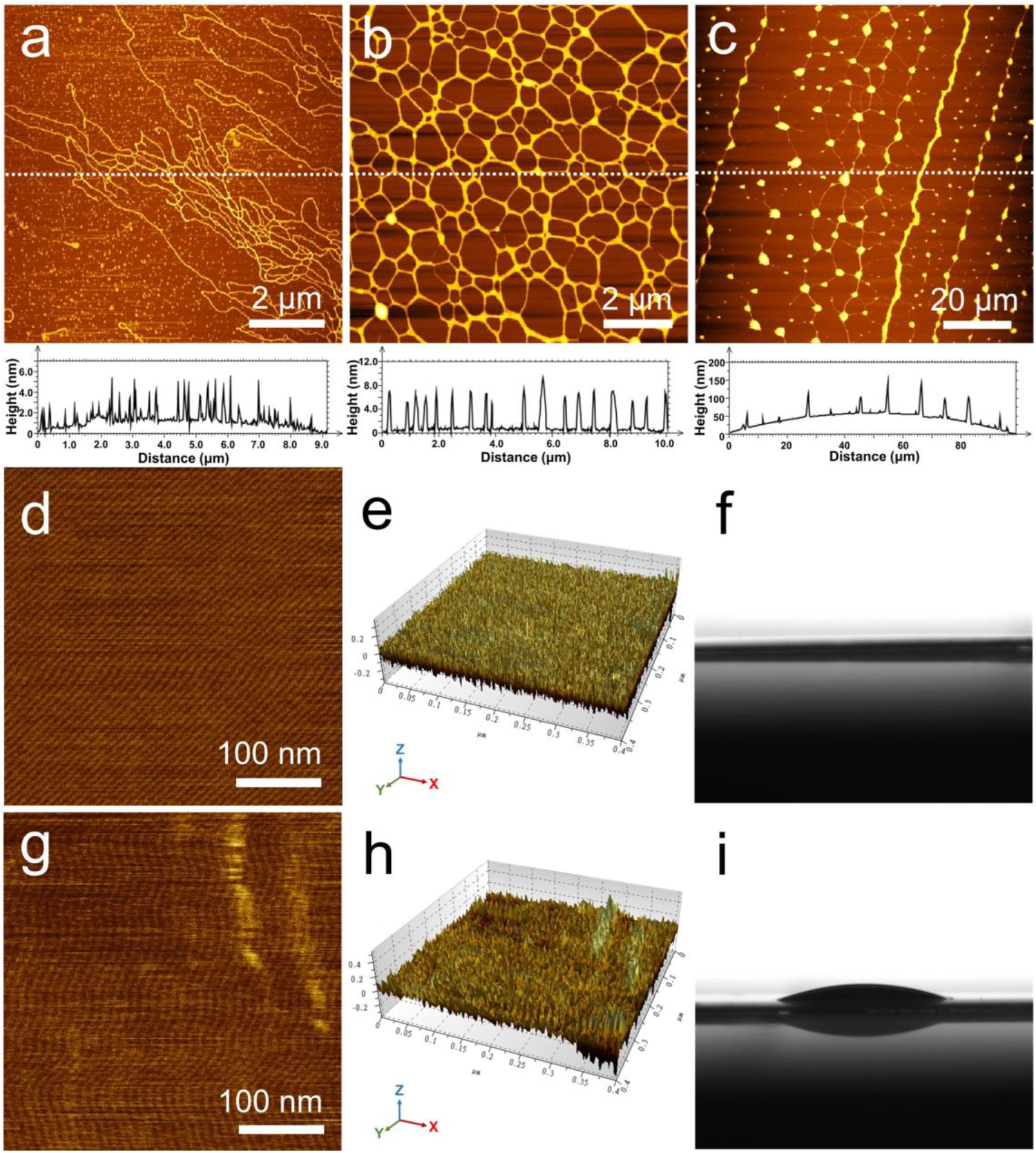
Typical AFM images and the corresponding height profiles of 10 ng/μL DNA on the Mg^2+^ modified mica produced under the different HEF DC voltages: 0 V (a), 0.5 V (b) and 1.0 V (c), respectively. The AFM images and the corresponding 3D topographic maps of bare (d, e) and Mg^2+^ modified (g, h) mica surfaces. Pictures (f) and (i) are the water droplet on the bare and Mg^2+^ modified mica surfaces, respectively.

For a detailed discussion, the DNA self-assembly was the result of multiple forces acting together including the electrostatic attraction and the hydrophilic force between DNA and the mica surface, the attraction force between DNA molecules (37) and the electric field force. The Mg^2+^ plays a role of bridge between the phosphate groups of DNA molecules and the surface of mica, which could enhance the electrostatic attraction and the hydrophilic force between the DNA and the mica surface. The AFM images as well as the corresponding 3D topographic maps of bare and Mg^2+^ modified mica surfaces are shown in Fig. 5d, Fig. 5e, Fig. 5g and Fig. 5h, indicating that the surface roughness has increased after the treatment of Mg^2+^ modification. The increased surface roughness on the one hand increased the contact opportunity between the DNA molecules and mica surface. On the other hand, from Fig. 5f and Fig. 5i, the increase of surface roughness led to the decrease of the surface hydrophilicity, thus the DNA solution was as a result not easy to spread on the mica surface, which contributed to the accumulation of DNA to a large extent under the application of electric field.

In addition to the self-assembly behavior of DNA in the horizontal electric field, the arrangement of DNA molecules in the vertical electric field was also investigated. Fig. 6 displays the typical AFM images of 1 ng/μL DNA on the bare mica prepared under different VEF DC voltages. Among them, the direction of the applied vertical electric field was upward for Figs. 6a~c and downward for Fig. 6d. It could be seen that the distribution of DNA molecules was obviously affected by the change of voltage. At a low voltage of 0.5 V, the DNA molecules were interconnected to form a relaxed DNA network showing a similar morphology to that in Fig. 3a. When the voltage increased to 0.75 V as shown in Fig. 6b, the density of DNA networks decreased and there became a tendency for the DNA molecules to be stretched. Moreover, the surface of the area around the DNA molecules seem to be rough owning to the deposition of residual buffer salt (38). Fig. 6c shows that at the voltage of 1.0 V, a higher amount of residual buffer salts deposited into a dense thin layer with a great deal of microporous structures on the mica surface, which is not found in the sample prepared in the horizontal electric field. The stretched DNA chains were embedded in the as-formed residual buffer salt layer and there was no distinct intermolecular contact, crossover and overlap between them.

**Fig. 6.**
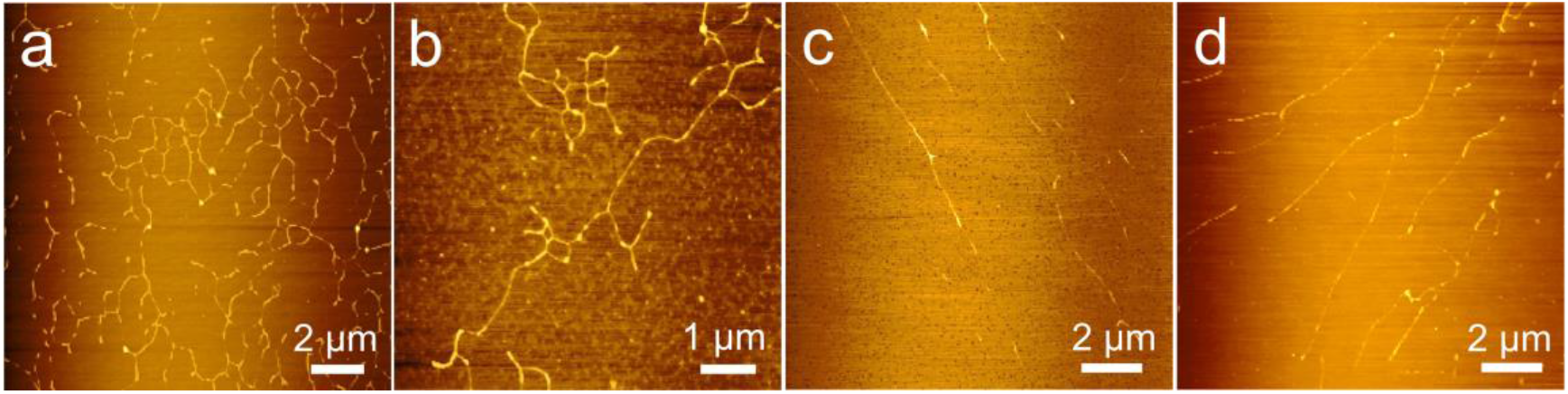
Typical AFM images of 1 ng/μL DNA on the bare mica produced under the vertical upward DC voltages of 0.5 V (a), 0.75 V (b), 1.0 V (c) and downward VEF DC voltage of 1.0 V (d).

A possible explanation for the above results could be that the higher electric density promoted the electrostatic attraction and the hydrophilic force between the DNA and the mica surface rather than the attraction force between the DNA molecules, which contributed to the enhanced stretching effect of DNA molecules with the increase of the voltage. Although the upward electric field at a relatively high voltage did make a difference on stretching DNA molecules, the formation of this residual buffer salt layer was not expected.

Correspondingly, for the downward vertical electric field at 1.0 V as shown in Fig. 6d, the DNA molecules exhibit stretching result similar to that shown in Fig. 6c, but there was no obvious buffer salt crystallization around the stretched DNA molecules, which indicated that a surface with rich positive charges is more conducive to the formation of residual buffer salt layer.

Fig. 7 shows the distribution of DNA molecules at different concentrations of DNA solution under the 1.0 V upward vertical electric field. For a more detailed examination of DNA with the concentration of 1 ng/μL, the DNA chain embedded in the residual buffer salt layer can be clearly seen from the local magnification of Fig. 7a. The height of the DNA chain was measured to be approximately 2.5 nm from the height map, which was about 5 dsDNA high, and the thickness of the residual buffer layer was nearly 1.0 nm. At the concentration of 5 ng/μL, the DNA molecules were accumulated to form parallel long arrays as shown in Fig. 7b. From the edge of the arrays, the tentacle-shaped DNA fibers stretched out and contacted with each other to form a network structure. There was no residual buffer layer formed on the mica surface. The DNA arrays mainly composed of two typical aggregation patterns, as shown in the local magnification maps. One structure of the patterns was formed by the tightly intertwining and stacking of DNA molecules, which contributed to a relatively higher height and narrow width of the arrays. At the edge of the arrays where the DNA chains stretched out, the thin DNA films with the height of about 3~4 nm could be found. For another one, a large amount of DNA molecules curled up under the action of electric field and accumulated into layered structure with the height of about 2 nm. Fig. 7c displays that as the concentration increased to 10 ng/μL, the regular morphology is not obtained owning to the serious agglomeration of DNA molecules.

**Fig. 7.**
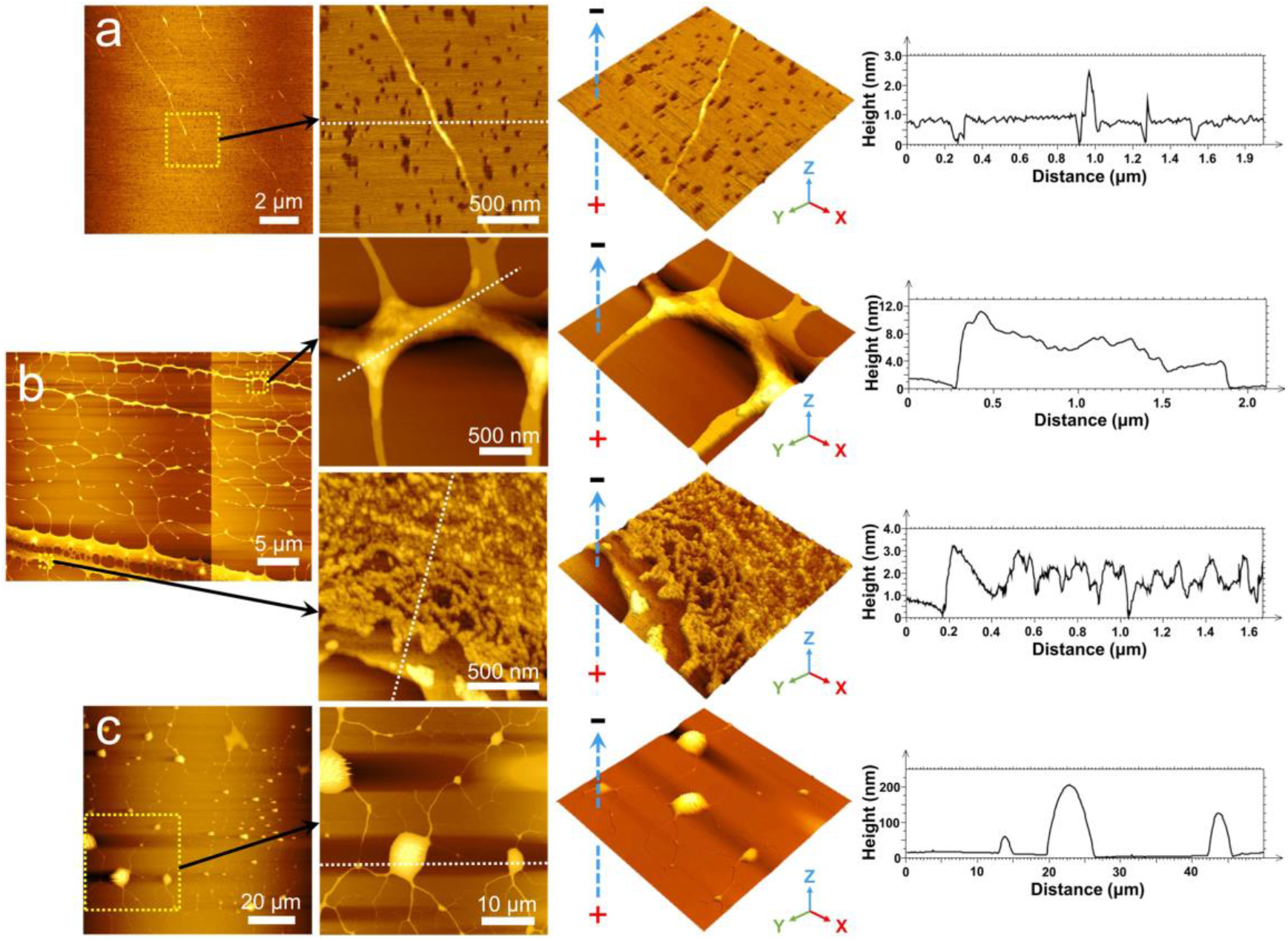
Typical AFM images (the first column), local amplification images (the second column) and the corresponding 3D topographic maps (the third column) of 1 ng/μL (a), 5 ng/μL (b) and 10 ng/μL (c) DNA on the bare mica produced under the 1.0 V VEF DC voltage. (Note: The blue dotted arrow on the 3D topographic maps represents the direction of the vertical electric field.) The fourth column is the height profiles of the transversal section marked with the white dashed line in the local amplification images.

## Conclusions

In summary, large-scale morphologically controlled self-assembled DNA networks were fabricated by the synergistic effect of DC electric field. By comparing the AFM images the samples under different electric field directions, it was shown that the horizontal electric field was more advantageous to the formation of DNA networks with more regular structures. Moreover, at the same DNA concentration, the change of horizontal electric field intensity did not affect the height of DNA network significantly. It was also found that the modification of Mg^2+^ increased the aggregation of DNA molecules under the electric field, and thus formed higher DNA networks. In addition to the formation of DNA self-assembly networks, obvious DNA molecular stretching results at both horizontal and vertical electric fields were obtained at low DNA concentrations. The above information may help to prepare controllable self-assembled DNA networks with diverse patterns and will also promote the application of DNA in the field of nanotechnology.

## Conflict of interest

The authors declare no conflicting financial interest.

## Author contributions

M.G. designed research, performed research, analyzed data, and wrote the article. J.H. designed research and analyzed data. Y.W. contributed analytic tools and analyzed data. M.L. contributed analytic tools and analyzed data. J.W. contributed analytic tools and analyzed data. Z.S. analyzed data. H.X. analyzed data. C.H. designed research, performed research, and wrote the article. Z.W. designed research, performed research, contributed analytic tools, and wrote the article.

## Acknowledgements

We are grateful to the National Key R&D Program of China (No.2017YFE0112100), EU H2020 Program (MNR4SCELL No.734174), Jilin Provincial Science and Technology Program (Nos. 20160623002TC, 20180414002GH, 20180414081GH, 20180520203JH and 20190702002GH), and “111” Project of China (D17017). This work was also partly supported by Changli Nano Biotechnology (China).

